# Allele-specific expression reveals interactions between genetic variation and environment

**DOI:** 10.1101/025874

**Authors:** David A. Knowles, Joe R. Davis, Anil Raj, Xiaowei Zhu, James B. Potash, Myrna M. Weissman, Jianxin Shi, Douglas F. Levinson, Sara Mostafavi, Stephen B. Montgomery, Alexis Battle

## Introduction

The impact of environment on human health is dramatic, with major risk factors including substance use^1^, diet^2^ and exercise^3^. However, identifying interactions between the environment and an individual’s genetic background (GxE) has been hampered by statistical and computational challenges^4,5^. By combining RNA sequencing of whole blood and extensive environmental annotations collected from 922 individuals^6^, we have evaluated GxE interactions at a cellular level. We have developed EAGLE, a hierarchical Bayesian model for identifying GxE interactions based on association between environment and allele-specific expression (ASE). EAGLE increases power by leveraging the controlled, within-sample comparison of environmental impact on different genetic backgrounds provided by ASE, while also taking into account technical covariates and over-dispersion of sequencing read counts. EAGLE identifies 35 GxE interactions, a substantial increase over standard GxE testing. Among EAGLE hits are variants that modulate response to smoking, exercise and blood pressure medication. Further, application of EAGLE identifies GxE interactions to infection response that replicate results reported *in vitro*^7^, demonstrating the power of EAGLE to accurately identify GxE candidates from large RNA sequencing studies.

## Main text

Phenotypic variation results from the combined effect of environment and individual genetic background. Many environmental and behavioral influences have been shown to substantially affect human disease risk^1,2,8^, and in model organisms gene-by-environment (GxE) interactions have been shown to be pervasive^9,10^. However, the prevalence and importance of GxE in human health is not well characterized, and identifying associations on a large scale in human populations has been challenging^4,5^. There are genetic variants that affect individual drug metabolism and response^11^, but only a few GxE interactions with disease have been identified^12,13^, with mixed results in replication^14^. Targeted experimental approaches are not always practical, and detection of GxE from genome-wide data faces considerations including small genetic effect sizes for most complex traits and high multiple hypothesis-testing burden.

In this study, we analyzed GxE in the context of transcriptomic phenotypes; cellular traits can reflect or even mediate disease risk, and the effects of genetic variation on gene expression are large enough for well-powered, genome-wide detection of expression quantitative trait loci (eQTLs) even in modestly-sized cohorts. Indeed, recent genetic studies of gene expression using RNA-sequencing have found thousands of eQTLs with high reproducibility^6,15–17^. Gene expression can also reveal the impact of environmental factors^18–21^, and recently, studies have begun to evaluate GxE interactions using transcriptomic data. *In vitro* immune stimulation has been used to detect hundreds of GxE interaction effects on gene expression in both human monocytes^7^ and dendritic cells^22,23^. Further, agnostic to the specific environment involved, the presence of extensive GxE interactions on the transcriptome is supported by variance eQTL mapping^24^ and allele specific expression^25^ in mono- and dizygotic twins. However, transcriptomic GxE mapping has not yet been performed for most major environmental risk factors. The emerging availability of cohorts with RNA-sequencing of primary tissue and well-curated clinical data provides an opportunity to test GxE interactions for specific environmental factors on a large scale. Still, significant technical challenges remain. Specifically, both biological and technical factors that vary across samples, such as batch effects and correlation among environmental factors, can confound the detection of GxE from transcriptomic data. Second, as discussed below, we observe that standard methods for testing GxE using gene expression are still underpowered, even in large cohorts.

To improve power to discover GxE interactions, we developed EAGLE (Environment-ASE through Generalized LinEar modeling), a novel method to test for GxE interactions using allele specific expression (ASE). Intuitively, observing that allelic imbalance of a gene associates with a particular environmental factor suggests that there is a *cis*-regulatory effect whose impact on expression is modulated by that environment. For example, an environmentally responsive transcription factor that binds to one allele better than to the other allele (Figure 1A) would result in allelic imbalance of the target gene in that environmental context. By comparing two alleles in the same sample, ASE provides an “internally matched” measure that inherently provides improved control for batch effects and other forms of confounding technical variation (Supplementary Figure S1). We designed EAGLE to use a binomial generalized linear mixed model (GLMM), predicting the relative number of RNA-seq reads from each allele at exonic, heterozygous loci under different environmental conditions. By directly modeling allelic read counts, rather than a simple continuous estimate of allelic imbalance, EAGLE improves power and additionally is able to model and account for over-dispersion inherent in RNA-seq data. As in previous analyses^26,27^ we have observed that allelic read counts display extra-binomial variation. While some apparent over-dispersion is likely to come from genetic and biological causes, PCR amplification and other technical factors may also contribute which when ignored lead to false positive associations (Supplementary Figure S2). EAGLE estimates a per-locus overdispersion parameter (random effect variance) that accounts for both technical overdispersion and extrinsic variation between individuals. Statistical power is shared across loci by learning a genome-wide prior on variance parameters. Since EAGLE is a generalized linear mixed model, it is straightforward to add additional covariates. In particular, we control environment-independent *cis*-eQTL by including an indicator variable denoting whether the lead eQTL (found by total expression analysis) for the gene is heterozygous. Similarly, EAGLE can be used to identify associations with other, non-environmental factors, such as the identification of aseQTLs (Supplementary Figure S3). EAGLE provides a flexible framework for modeling influence of both technical and biological factors on ASE while accounting for extra-binomial variation in sequencing data.

**Figure 1:**
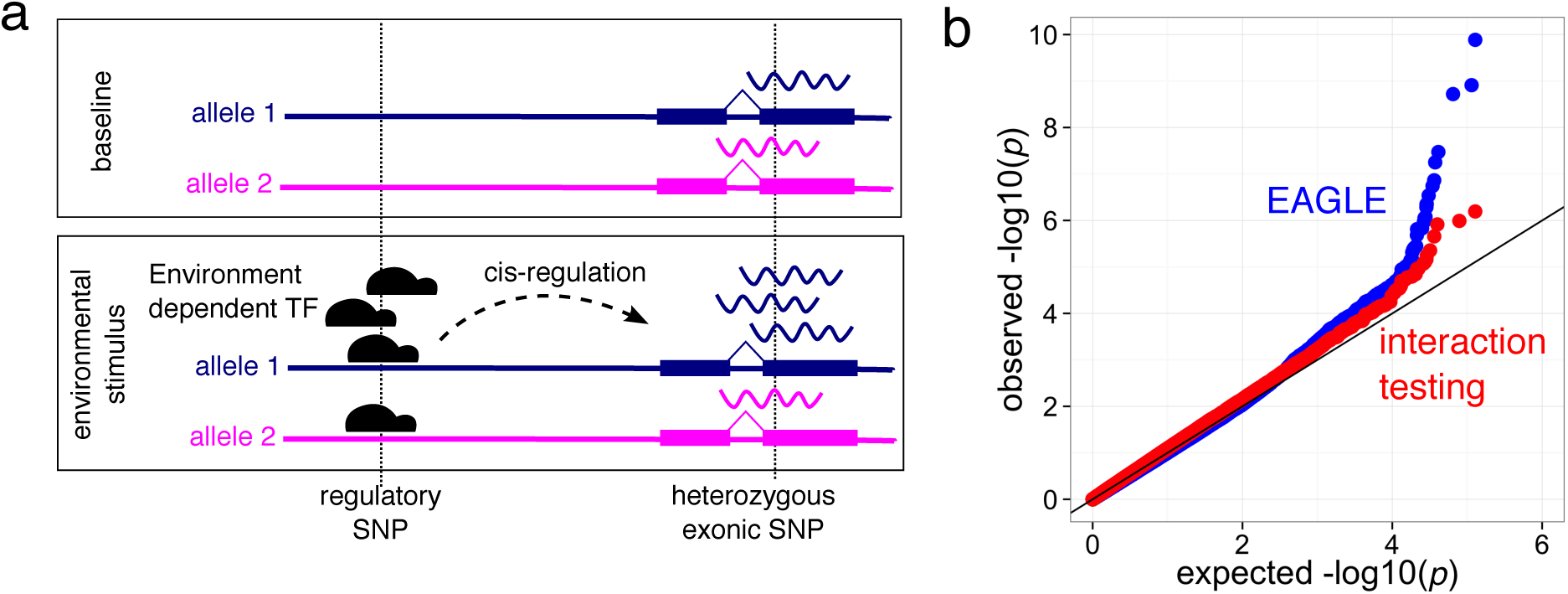
EAGLE associates allelic specific expression (ASE) with environmental covariates to detect GxE interactions. (**a**) Allelic imbalance can be driven by allele specific binding of an environmentally responsive transcription factor. (**b**) Using ASE increases power relative to standard interaction testing in the DGN cohort across 30 environmental variables. EAGLE provides an internally controlled test and integrates across the *cis*-regulatory landscape of a gene.

We applied EAGLE to the discovery of GxE interactions from a large publicly-available cohort of 922 individuals with RNA-sequencing data from the Depression Genes and Networks study^6^. This study has high power to detect eQTLs, with 79% of tested transcripts having an eQTL for total expression at a conservative FDR (5%). In addition, diverse annotations are available describing medication use, behavior, and other environmental factors for each individual. The samples come from a primary tissue, enabling accurate analysis of environmental influences on the transcriptome; indeed, we detect thousands of environmentally responsive genes (Supplementary Figure S4).

We tested for EAGLE associations between 30 environmental factors (Supplementary Table S1) and ASE of 8795 genes (Methods). We found 35 significant associations at an FDR of 10% (Supplementary Table S2). Among these, we detected a GxE interaction between exercise before blood draw and *DYSF*, a skeletal muscle repair protein. Mutations in *DYSF* cause the recessive muscular dystrophy *dysferlinopathy,* with progression of the disease being exercise level dependent^28^, indicating a disease relevant GxE interaction for this gene. We further detected a GxE interaction for blood pressure medication with *NPRL3*, part of the *NPR3* protein family involved in homeostasis of fluid volume (Figure 2a). We also observed that higher BMI is associated with increased allelic imbalance of *VNN1*, which is associated with high-density lipoprotein cholesterol^29^, prevents lipid peroxidation^30^ and is predicted to be causally related to omental fat pad mass^31^.

**Figure 2:**
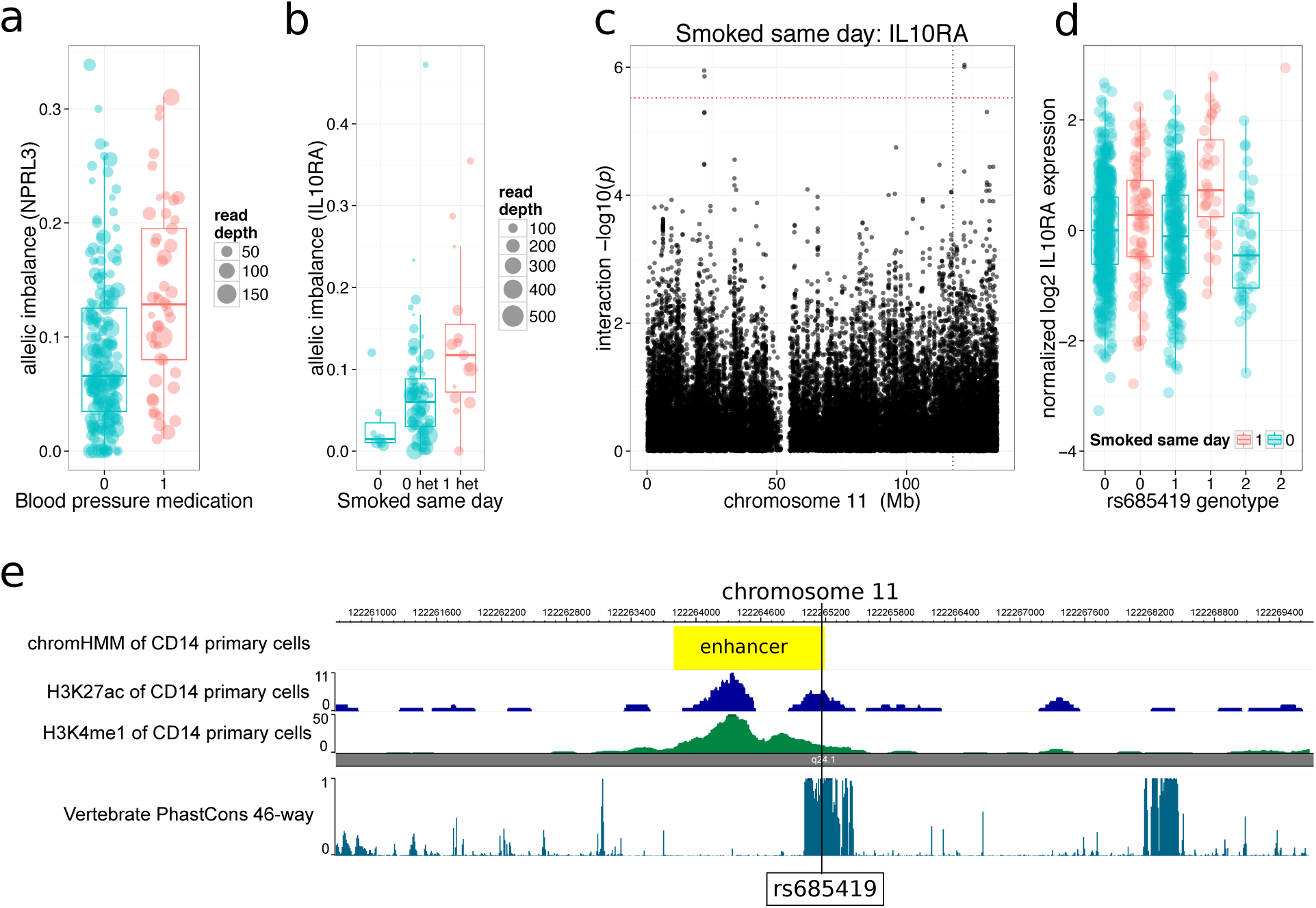
EAGLE detects GxE interactions missed by standard interaction QTL testing. (**a**) Blood pressure medication modulates regulation of *NPRL3*, involved in fluid homeostasis. (**b**) Smoking interacts with regulation of *IL10RA* (*interleukin 10 receptor-α*). (**c-e**) Using standard interaction QTL testing as a second phase within EAGLE hits, we detect *rs685419* as a promising candidate variant for smoking association with IL10RA, lying 4Mb from the TSS in a conserved region corresponding to an enhancer in CD14+ primary cells.

As a baseline against which to benchmark EAGLE’s power, we also detected GxE interactions on total expression using a standard linear model interaction test (Methods). Using Bonferroni correction per gene, since there is no appropriate permutation strategy for interaction testing^32^, followed by controlling the FDR at 10% we find only four associations across the 30 tested environmental factors. Thus, EAGLE shows much greater power to detect GxE interactions than standard interaction QTL testing (Figure 1B). Results from EAGLE or standard methods could represent interactions with (potentially unmeasured) factors that are correlated with the tested environmental variables. EAGLE however should be less susceptible to false positives from some technical confounders (Supplementary Figure S1). Overall, the improved power may derive from multiple sources, including the controlled, within-individual nature of our ASE-based test along with the direct modeling of read counts. Further, EAGLE implicitly integrates over the entire *cis*-regulatory landscape of a gene rather than explicitly testing a specific candidate SNP, reducing the multiple hypothesis-testing burden and potentially captures the contribution of multiple regulatory variants.

EAGLE does not directly test individual candidate SNPs responsible for the association between environment and ASE. However, we applied a two-step procedure based on EAGLE for finding candidate variants driving GxE associations that still yields more hits than standard interaction QTL testing. In the first step, EAGLE was used with a lenient FDR of 0.2 to give a shortlist of 57 environment-gene associations. In the second step, we looked for candidate variants, within 1Mb of the TSS, using EAGLE combined, through meta-analysis, with standard interaction testing (see Methods). For 15 out of 57 associations we found a *cis*-SNP with a nominally significant interaction QTL after conservative Bonferroni correction across tested SNPs (*p*<0.05; Supplementary Table S3). Those with no candidate variant hit may arise from variants outside of the 1MB window, rare variants, or non-genetic factors. Some SNPs were not testable using EAGLE because not enough double heterozygous individuals were available. In this case, we used standard interaction testing alone. For the association between *smoked same day* and *IL10RA* (Benjamini-Hochberg *q=0.13*, see Figure 2b) the top candidate variant (*p* = 9 ×10^−7^) is *rs685419* which lies 4Mb from the TSS of *IL10RA* (*interleukin 10 receptor-α*) in a conserved CD14 primary cell enhancer (Figure 2c-d). Polymorphisms in *IL10* itself have been associated with the rate of lung function decline in firefighters^33^. Since many diseases result from the combined effects of genetics and environment we investigated whether any of our candidate GxE variants, or variants in linkage disequilibrium (LD), are known genetic risk factors for disease using the NHGRI-EBI GWAS (accessed 6/17/2015)^34^ and Immunobase (available at www.immunobase.org; accessed 6/21/2015) catalogs. We identified eight disease-associated variants (Supplementary Table S4). For example, we found that *rs1538257*, which is the top candidate variant to modulate BMI’s association with *LGALS3* expression, is in LD (R^2^=0.55) with *rs2274273*, which is associated with *LGALS3* protein levels (*p* = *2* ×10^−188^). Interestingly, in mice, *LGALS3* has been shown to have a protective role in obesity induced inflammation and diabetes^35^.

Next, we sought to characterize the properties of the genes whose genetic regulation is modulated by each environment. Since the number of genome-wide significant associations remains relatively modest even with the improved power from EAGLE, we performed enrichment analysis using the top 50 associations for each environment. We first tested these associations against a curated set of pathways taken from GO, KEGG and BioCarta (restricted to those with fewer than 100 genes), using a standard hypergeometric test with the entire set of genes tested by EAGLE as the background. The strongest enrichment is for smoking and the *BioCarta CCR5 pathway*. *CCR5* itself has been implicated in smoking induced emphysema^36^. Since our hypothesis is that GxE interactions for gene expression are often driven by allele specific binding of environmentally-responsive transcription factors, we tested for enrichment of transcription factor binding sites (TFBS) proximal to environment-associated genes. We used the union of TFBSs detected by CENTIPEDE^37^ from *DNase I* hypersensitivity data for seven blood cell types^38^ (see Supplementary Material Section 4). Since we only expect to see GxE when there is corresponding genetic variation, we filtered for TFBS within 5kb of each gene that also contained at least one variant previously identified in the 1000 Genomes Project, resulting in an average of 7.7 TFBS per gene across 282 TF motifs. We again used a hypergeometric test for enrichment for each environment (Figure 3B). The strongest association (p=10^-4^) is for smoking-associated genes and the transcription factor *TBX4*. *TBX4* is known to be regulated by *SOX9*, variants in which influence lung function specifically in smokers^39^. Additionally, genes showing a GxE interaction for blood pressure medication are enriched in binding of *SP1*, which is known to respond to antihypertensive drugs^40^ and regulates angiotensin receptor transcription^41,42^.

**Figure 3:**
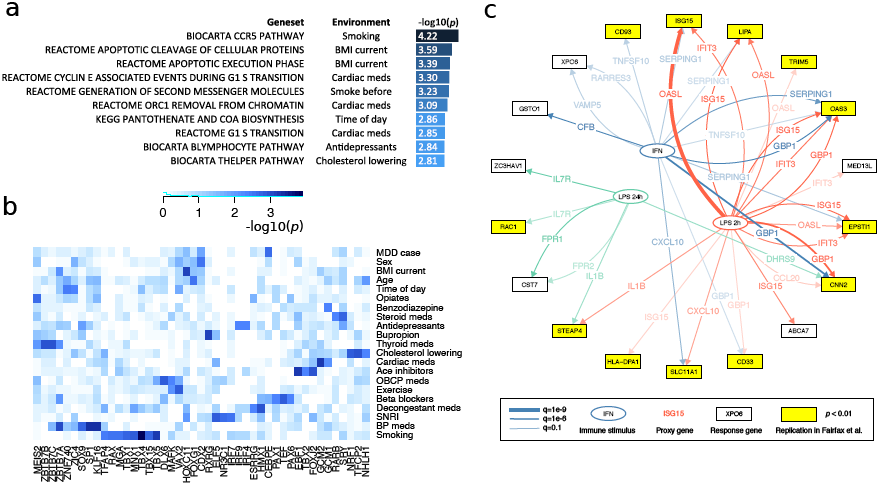
Pathway and transcription factor binding site enrichment of genes with GxE interactions for each environment reveals shared regulation. Uncorrected *p*-values are shown. (**a**) The strongest associations between the top 50 genes for each environment and GO, KEGG and BioCarta pathways with fewer than 100 genes. (**b**) Enrichment of CENTIPEDE predicted TFBS within 5kb of the TSS of the top 50 genes associated with each environment. (**c**) EAGLE recapitulates GxE interactions discovered using immune stimulation of monocytes in vitro. We used genes differentially expressed under immune stimulation in vitro as proxies for the environment (stimulus). The genes detected by EAGLE as being modulated by these environmental proxies replicate in the in vitro data: i.e. they have detectable response QTLs. Network depicts all EAGLE predictions for each stimulus, with replicating interactions highlighted in yellow; each edge is annotated with the tested proxy gene for reference.

Further, we investigated additional evidence for co-regulation of EAGLE hits of each environment based on *trans*-eQTLs. DGN’s relatively large sample size enables the detection of inter-chromosomal *trans*-eQTLs (138 unique trans-eQTL genes at an FDR of 0.05 ^6^). Applying a relaxed, nominal *p*-value threshold of 10^-5^ yielded a trans-network with 55,313 edges involving 48,163 SNPs and 7473 genes. We investigated whether the top 50 genes associated by EAGLE for each environment tend to share distal regulatory SNPs in this network. Against an empirical null distribution generated by randomly sampling sets of 50 genes from those tested for each environment, we found the number of SNPs regulating more than one of the 50 genes is significantly increased (*p*<0.05) for age, exercise, family history of depression and opiate use. The trans-network involving SNPs regulating more than one gene in the top 50 list for exercise is shown in Supplementary Figure S5. Interestingly all five of the genes (*IFIT2*, *MX2*, *IFI44L*, *ADAR*, *RSAD2*) implicated in this network are interferon inducible, highlighting the impact of exercise on immune response^43^.

We investigated the degree to which EAGLE analyses, conducted within a large cohort, recapitulate GxE interactions discovered *in vitro*. Specifically, the interplay of immune stimulation, gene expression and genetics has been characterized in several recent *in vitro* studies: Barreiro *et al.* infected primary dendritic cells (DCs) with *Mycobacterium tuberculosis*^23^, Lee *et al.* stimulated DCs with lipopolysaccharide (LPS), influenza virus, or *IFN-β*^*22*^, and Fairfax *et al.* exposed CD14*+* monocytes to interferon-γ (IFN-γ) and LPS for 2 or 24 hours^7^. All three of these studies found more eQTLs under stimulated conditions than in steady state, and discovered corresponding GxE interactions. To test if these interactions are detectable in our cohort, we focused on the Fairfax *et al.* study^7^ due to its large sample size, genome wide transcriptomic profiling and choice of interferon-γ (IFN-γ) and LPS immune stimulation (likely to be relevant in a population sample). Direct measurements of infection and immune activity are not available for the DGN cohort. We therefore used the expression levels of the top differentially expressed genes for each stimulus as “proxies” for the environment. Specifically, we identified 25, 16, and 26 genes, for LPS at 2h, LPS at 24h and IFN-γ respectively, with an absolute log-fold change greater than 4 in the Fairfax *et al.* data. We then applied EAGLE genome-wide to find association between ASE and gene expression levels for each proxy gene. We exclude tests for interactions between proxy genes and allelic balance of genes on the same chromosome since such an association could represent direct *cis*-regulation rather than an interaction. At an FDR of 10%, we found 26, 6 and 14 GxE interactions for LPS at 2h, LPS at 24h and IFN-γ respectively. To test whether these interactions were also detected in Fairfax *et al*. we compared the reported *t*-statistics for the lead eQTL under the naïve and stimulated condition (Supplementary Material Section 5, Supplementary Figures S7-8). At a nominal *p*-value threshold of 10^-4^ we found that 11/26, 3/6 and 6/14 interactions replicated for the three stimuli respectively (Figure 3c). To assess the significance of this replication, we generated an empirical null distribution using randomly chosen sets of environmental proxy genes not differentially expressed in response to any of the Fairfax *et al.* stimuli. This analysis gave empirical *p*-values for the observed replication of 0.048, 0.06, and 0.029 respectively, or 0.0017 for the overall replication frequency.

The results obtained by applying EAGLE to the DGN cohort demonstrate that careful analysis of allele specific expression from RNA-seq is an effective way to identify candidate interactions between genotype and environment that contribute to transcriptional variation. A key finding is that by modeling allele-specific read counts directly, EAGLE offers significantly improved power to detect GxE interactions over standard linear modeling of total gene expression. This is consistent with observations from recent studies of ASE in contexts other than GxE, such as QTL analysis^44,45^. The associations and variants detected by EAGLE indicate that common environmental risk factors, including substance use, exercise and BMI do in fact interact with individual genetic variation in regulation of gene expression. We also report a number of associations with potential consequences on disease risk. Despite the large increase in power, the overall number of associations remains modest, with 35 detected for 30 environments from a sample of 922 individuals, indicating that GxE effects on gene expression are not prevalent with large effect sizes compared with additive effects. Additionally, there are allele-specific, *cis*-regulatory mechanisms other than genetic effects that could potentially explain some of the associations discovered, for example epigenetic regulation of expression. Finally, we note that the DGN samples analyzed here are from whole blood, which may mask GxE effects limited to a specific cell-type, although current multi-tissue eQTL studies indicate that *cis*-eQTLs are generally highly shared across tissues^16,46^. In conclusion, despite the challenges in analyzing GxE interactions, we show it is possible to leverage the novel information provided by large RNA-seq cohorts to unravel the modulation of genetic effects by environmental factors relevant to human disease.

EAGLE offers an extensible framework for robust detection of factors contributing to allelic imbalance across samples, and may be applied in various settings, such as the detection of aseQTLs (Supplementary Figure S3) or the reconstruction of regulatory networks. EAGLE provides a general method for accounting for over-dispersion and for modeling effects of technical covariates on both mean and variance of ASE. For instance, EAGLE could also be applied to *in-vitro* studies of GxE where RNA-seq is available for both control and perturbed conditions for the same cell-line or individual, with minor modification to the GLMM to account for correlated measurements. Additionally, the use of “proxy genes” to represent unmeasured environmental factors opens up a number of applications on a large scale with existing data and at comparably low cost, as demonstrated by our replication of infection response QTLs from Fairfax *et al.*^7^. As RNA-seq datasets become widely available, we envisage that EAGLE will be appropriate to obtain additional power to detect individual differences in environmental response for a wide range of contexts and studies. More generally, EAGLE is a useful tool for understanding the combined effects of external stimuli, genetic variation, and cellular networks on regulation of gene expression.

## Methods

### Interaction QTL testing

Total expression was quantified as previously described^6^, including controlling for known and latent confounders using HCP^47^. We quantile normalize each gene to a standard normal distribution to remove outliers, and perform standard interaction testing to find GxE effects for the 8795 genes testable using ASE. For a specific combination of SNP, gene and environment consider the null model *H*_*0*_ and alternative model *H*_1_.

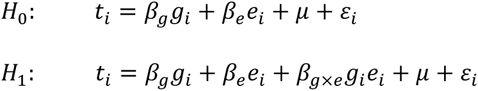

where *t*_*i*_ is normalized total expression for individual *i*, *g*_*i*_ is the genotype of the SNP encoded as {0,1,*2*}, *e*_*i*_ is the environmental factor, *β*_*g*_, *β*_*e*_, *β*_*g × e*_ are genetic, environment and interaction effect sizes respectively and μ is an intercept. Under the null the likelihood ratio 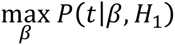 is *χ*^*2*^-distributed with one degree of freedom, which allows us to obtain a well calibrated *p*-value. We test all SNPs within 200kb of the TSS (obtained from GENCODE, release 20). Since there is no appropriate permutation strategy for testing interaction terms ^32^, we were constrained to using Bonferroni correction to obtain an approximate gene level *p*-value. The gene level *p*-values for a particular environment are then adjusted using the Benjamini-Hochberg procedure to control the FDR at a pre-specified level.

### EAGLE model

We first present the model itself and then motivate the various modeling choices. The null model *H*_*0*_ is

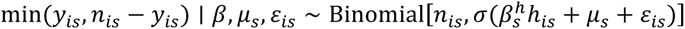

and the alternative model *H*_1_ is

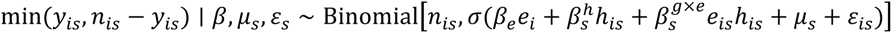

where *y*_*is*_ is the alternative read count for individual *i* at locus *s*, *n*_*is*_is the total read count, σ(x) = 1/(1 + e^−*x*^) is the logistic function, *h*_*is*_ denotes whether the top *cis*-eQTL is heterozygous, *μ*_*s*_ is an intercept term to take into account unexplained allelic imbalance unrelated to the environment and *∈*_*is*_|v ∼ N(0, v_s_) is a per individual per locus random effect modeling overdispersion. This model can be derived by assuming the log expression of each allele is linear in the environment and SNP genotype (see Supplementary Material Section 1. The variance itself is given an inverse gamma prior *IG*(*a, b*). We learn the hyperparameters *a*,*b* across all genes. We expect that environmental effects on ASE are usually mediated by one or more causal *cis*-regulatory genetic variants, which would often be in linkage disequilibrium with the locus where ASE is measured. However, some responsive individuals may have different causal sites and therefore may exhibit opposite direction of allelic effect. EAGLE gains power by testing just a single association statistic per gene, rather than modeling each possible causal site and incurring a large multiple testing burden, but therefore cannot assume a consistent direction of allelic effect across the cohort. Additionally, linkage disequilibrium may be weak, especially for more distal elements. The EAGLE model is applicable in settings where causal sites vary between individual and also handles unphased data. We model the absolute deviation from allelic balance by considering min (*y*_*is*_, *n*_*is*_ - *y*_*is*_) rather than the minor allele count *y*_*is*_ itself. This is analogous to using 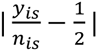 as a quantitative measure of allelic imbalance, but maintains the count nature of the data. We also experimented with introducing explicit auxiliary “flipping” variables to provide implicit phasing, but found this was susceptible to over-fitting.

### Accounting for cis-regulation

Standard cis-eQTL analysis allowed us to identify proximal genetic variants associated to the expression of each gene. These variants often explain a significant proportion of observed ASE. To account for this, we add a dependence on *h*_*is*_, an indicator of whether the top *cis*-eQTL for the gene containing locus *s* is heterozygous in individual *i*. Additionally, in some cases one of the known *cis*-eQTLs could be the variant through which the environment influences the observed ASE, which we model by including an interaction term *h*_*is*_*e*_*is*_ (see Supplementary Material Section 2 for further details). We approximately integrate over the random effects ϵ_*is*_ and per locus variance v_*s*_ using non-conjugate variational message passing^48^ while optimizing the coefficients *β* and hyperparameters *a, b* (Supplementary Material Section 3).

### Parameter estimation and inference

Holding the overdispersion hyperparameters *a*,*b* fixed we fit both the alternative and null models at each locus and use the variational lower bound as an approximation to the true marginal likelihood for each model, allowing us to calculate an approximate likelihood ratio. It is not obvious that the usual asymptotic theory should hold here since a) our data is not normally distributed, b) we only have an approximation of the true likelihood, and c) our model incorporates random effects terms. To investigate this we performed permutation experiments, using the conveniently valid strategy of separately permuting the individuals heterozygous or homozygous for the top *cis*-SNP^32^. These experiments show that our approximate likelihood ratios do in fact follow the asymptotic *χ*^2^ distribution quite closely, while being slightly conservative (see Supplementary Figure S6). Therefore we choose to use the nominal likelihood ratio test *p*-values, avoiding having to run computationally expensive permutation analysis for every tested association.

### Data Access

Genotype, raw RNA-seq, quantified expression, covariates and environmental data for the DGN cohort are available by application through the NIMH Center for Collaborative Genomic Studies on Mental Disorders. Instructions for requesting access to data can be found at https://www.nimhgenetics.org/access_data_biomaterial.php, and inquiries should reference the “Depression Genes and Networks study (D. Levinson, PI)”.

### Software

EAGLE was developed in C++ and R 3.1.2 using RcppEigen and is available as an R package at https://github.com/davidaknowles/eagle.

## Acknowledgements

We would like to thank Jeff Leek for helpful comments. AB and SBM are supported by NIH R01MH101814. AB is supported by NIH R01 MH101820. SBM is supported by the Edward Mallinckrodt Jr. Foundation.

